# Exploring the phenotypic space and the evolutionary history of a natural mutation in *Drosophila melanogaster*

**DOI:** 10.1101/010918

**Authors:** Anna Ullastres, Natalia Petit, Josefa González

**Affiliations:** Institute of Evolutionary Biology (CSIC-Universitat Pompeu Fabra) Passeig Maritim de la Barceloneta 37-40. 08003 Barcelona, Spain

## Abstract

A major challenge of modern Biology is elucidating the functional consequences of natural mutations. While we have a good understanding of the effects of lab-induced mutations on the molecular- and organismal-level phenotypes, the study of natural mutations has lagged behind. In this work, we explore the phenotypic space and the evolutionary history of a previously identified adaptive transposable element insertion. We first combined several tests that capture different signatures of selection to show that there is evidence of positive selection in the regions flanking *FBti0019386* insertion. We then explored several phenotypes related to known phenotypic effects of nearby genes, and having plausible connections to fitness variation in nature. We found that flies with *FBti0019386* insertion had a shorter developmental time and were more sensitive to stress, which are likely to be the adaptive effect and the cost of selection of this mutation, respectively. Interestingly, these phenotypic effects are not consistent with a role of *FBti0019386* in temperate adaptation as has been previously suggested. Indeed, a global analysis of the population frequency of *FBti0019386* showed that clinal frequency patterns are found in North America and Australia but not in Europe. Finally, we showed that *FBti0019386* is associated with down-regulation of *sra* most likely because it induces the formation of heterochromatin by recruiting HP1a protein. Overall, our integrative approach allowed us to shed light on the evolutionary history, the relevant fitness effects and the likely molecular mechanisms of an adaptive mutation and highlights the complexity of natural genetic variants.

## INTRODUCTION

Understanding the functional consequences of naturally occurring mutations remains a largely open question in Biology. Most of our knowledge on the effect of mutations comes from the analyses of laboratory-induced mutations. However, it is not clear whether laboratory mutations are representative of mutations that arise and persist in natural populations (Kolaczkowski, et al. 2011; Rose, et al. 2011). First, most laboratory mutations studied are loss-of-function mutations that are most likely rare in natural populations and/or their effects are masked by the presence of buffering mechanisms (Landry and Rifkin 2012). Additionally, laboratory-induced mutations tend to be highly pleiotropic and it is difficult to infer which of the phenotypes might be targets of selection in nature (Kolaczkowski, et al. 2011).

The recent explosion in the number of studies aimed at identifying natural adaptive mutations in several organisms allow us to study the effect of natural genetic variants at an unprecedented scale (Gonzalez, et al. 2008; Turner, et al. 2010; Jones, et al. 2012; Huang, et al. 2014; Tobler, et al. 2014). These studies are revealing that mapping genotype to phenotype is even more complex than previously thought due to the prevalence of gene-by-environment, gene-by-gene interactions, and pleiotropy (Rockman 2012; Lehner 2013; Mackay 2014). First, being able to map a putatively adaptive mutation to its relevant phenotypic effect depends partly on finding the particular environmental conditions in which the mutation is adaptive (Paaby and Schmidt 2008; Storz and Wheat 2010). Taking into account environmental information of the geographical populations where putative adaptive mutations are identified, should thus help in mapping the mutation to its relevant phenotype. Second, epistatic interactions also affect the phenotypic outcome of mutations. The phenotypic effect of mutations could be enhanced or suppressed depending on the background being analyzed (Huang, et al. 2012). Additionally, several backgrounds should be analyzed to discard the effect of background mutations and reliably attribute the identified phenotypic effect to the candidate mutation (Burnett, et al. 2011). Third, many genes are linked to several traits (Paaby and Rockman 2013). In some cases, mutations can have antagonistic effects, *i.e.* beneficial effects in a trait/environment and deleterious effects on a different trait/environment. Pleiotropic mutations can also have beneficial effects on two different traits (McGee, et al. 2014). Tradeoffs are prevalent when selection acts on a single trait, while payoffs arise when multiple traits are selected for simultaneously (McGee, et al. 2014). Thus, if we want to fully characterize the effects of a given natural mutation, several phenotypes need to be studied (Mackay 2010; Guio, et al. 2014).

Finally, a comprehensive understanding of adaptation goes beyond identifying fitness consequences of adaptive mutations. Pinpointing the molecular mechanisms underlying adaptation is needed to provide conclusive support for the adaptive role of the mutation (Storz and Wheat 2010). Additionally, elucidating the evolutionary history of adaptive variation for fitness traits allows to start answering long-standing questions on the genetic basis of adaptation (Orr 2005).

In this work, we focused on characterizing the functional effects, the molecular mechanisms and the evolutionary history of a natural Transposable Element (TE)-induced mutation in *Drosophila melanogaster*: *FBti0019386* belonging to the *invader4* retrotransposon family (Gonzalez, et al. 2008; Gonzalez, et al. 2010; St Pierre, et al. 2014)*. FBti0019386* has been identified as a candidate adaptive TE insertion based on its population dynamics (Gonzalez, et al. 2008). González et al (2010) further showed that *FBti0019386* shows parallel clinal frequency patterns in North America and Australia suggesting that it is involved in adaptation to temperate environments. *FBti0019386* is inserted in the 5’UTR intron of *sarah* (*sra*) and 2.5 kb upstream of *Bicoid-interacting protein 1* (*Bin1*) in the 3R chromosomal arm (St Pierre, et al. 2014). *sra* laboratory mutants affect several biological processes such as egg activation, female meiosis, and long-term memory among others (Ejima, et al. 2001; Chang, et al. 2003; Ejima, et al. 2004; Horner, et al. 2006; Takeo, et al. 2006; Sakai and Aigaki 2010; Nakai, et al. 2011). In most cases, these phenotypes are the result of the deregulation of *calcineurin*, which is inhibited by *sra* (Takeo, et al. 2006; Sakai and Aigaki 2010; Nakai, et al. 2011). Laboratory-induced mutations affecting *Bin1*, a highly conserved transcriptional co-repressor, play a role during environmental stress response in *Arabidopsis* (Song and Galbraith 2006) and in *Drosophila* (Costa, et al. 2011). Thus, to identify the phenotypic consequences of *FBti0019386* mutation, we explored several candidate phenotypes previously associated with *sra* and *Bin1* mutants in different developmental stages, in different environmental conditions, and in flies with different genetic backgrounds.

Our results showed that *FBti0019386* increased in frequency in out-of-Africa populations due to positive selection and is associated with shorter developmental time and increased sensitivity to cold-stress. These two phenotypic effects together with the lack of clinal frequency patterns in Europe suggest that *FBti0019386* is not a clinal mutation. Finally, we also provide a mechanistic explanation for the effect of *FBti0019386* insertion: flies with *FBti0019386* insertion are associated with *sra* down-regulation most likely due to piRNA-mediated heterochromatin assembly.

## RESULTS

### *FBti0019386* flanking regions show signatures of positive selection

We tested whether the region flanking *FBti0019386* showed signals of positive selection (see Material and Methods). We found an extreme decrease of nucleotide diversity (π) in strains with *FBti0019386* insertion compared to strains without the insertion, which was accompanied by a decrease in Tajima´s D statistic (Table 1, Figure S1A and S1B and Table S1). The CL statistic, specifically designed to detect selective sweeps (Nielsen, et al. 2005), was also higher in flies with *FBti0019386* insertion compared to flies without the insertion, as expected if flies with the insertion show signatures of a selective sweep in the region analyzed (Table 1). We confirmed using simulations that values of nucleotide diversity, Tajima´s D, and CL statistic are statistically different from neutral expectations in flies with *FBti0019386* insertion but not in flies without the insertion (Table 1 and Table S2). To further test the significance of these results, we estimated the three statistics in random samples of the strains (see Material and Methods). None of the randomized datasets had lower nucleotide diversity, lower Tajima´s D, or higher CL value compared to the dataset of strains with *FBti0019386* insertion (Table 1, Table S3). Finally, we performed the Composite Likelihood Ratio (CLR) test comparing strains with and without the *FBti0019386*, and we found that it was significant: CLR= 24.40 p-value = 7.82x10^-7^. Moreover, this CLR value is bigger than any of the CLR values calculated in a random sample of 1,000 1kb-long regions from 3R chromosome, where *FBti0019386* is located (Table S4). Note that estimates of nucleotide diversity and Tajima´s D in these 1,000 regions also showed that these two statistics did not significantly differ between strains with and without *FBti0019386* insertion (Figure S1C and S1D). Overall, we found evidence of positive selection in the region flanking *FBti0019386* insertion suggesting that *FBti0019386* is an adaptive insertion.

**Table 1.** Summary of the analyses showing evidence of positive selection in the 1 kb region around *FBti0019386* insertion.

|  | Observed |  | Neutral simulations<sup>c</sup> |  |  |  |  |  | Re-sampling of strains |  |  |
| --- | --- | --- | --- | --- | --- | --- | --- | --- | --- | --- | --- |
|  |  |  | Mean |  | CI 95% |  | P-value |  | Mean | CI 95% | P-value |
|  | P<sup>a</sup> | A<sup>b</sup> | P | A | P | A | P | A |  |  |  |
| **Nucleotide diversity ($\pi$)** | 0.43 | 4.51 | 3.92 | 4.20 | 1.32, 7.81 | 1.33, 8.04 | p<0.001 | >0.05 | 3.35 | 2.78, 3.87 | p<0.001 |
| **Tajima's D** | -1.77 | 0.68 | -0.11 | -0.04 | -1.46, 1.62 | -1.41, 1.64 | 0.007 | >0.05 | 0.4 | -0.19, 1.02 | p<0.001 |
| **CL (log)** | -5.95 | -18.15 | -18.69 | -15.20 | -29.67, -8.80 | -25.89, -6.82 | 0.006 | >0.05 | -12.18 | -15.23, -8.81 | p<0.001 |

<sup>a</sup>P: dataset of lines with the insertion. <sup>b</sup>A: dataset of lines without the insertion. <sup>c</sup>Neutral simulations were performed with MS program using the parameter theta=4. For simulations with theta=5 please see supplementary Table S2.

### Exploring the fitness space of *FBti0019386*

To explore the phenotypic space of *FBti0019386* insertion, we investigated several traits related to the phenotypic effects of nearby genes: fecundity and egg hatchability associated with *sarah* mutant alleles. Related to egg hatchability, we investigated egg hatching time, egg-to-adult viability and developmental time. Additionally, we investigated cold stress, osmotic stress, and starvation stress since *Bin1* mutants have been shown to play a role in stress resistance.

### *FBti0019386* insertion does not affect fecundity or egg hatching

Laboratory mutant flies in which *sra* is under-expressed lay less eggs than wild-type flies and most of them do not hatch (Horner, et al. 2006). To check whether *FBti0019386* insertion has an effect on fecundity, we compared the number of eggs laid per female in outbred populations with and without the insertion (see Material and Methods). Our results showed that, on average, flies without the insertion laid slightly more eggs than flies with the insertion (t test, p-value = 0.047) (Figure 1A). However, the size effect of the mutation was not significant (Table 2).

**Table 2:** Odds ratios (OR) and confidence intervals (CI) for phenotypic experiments in flies with and without *FBti0019386*. N/A (see Material and Methods).

| Experiment | Strain | OR (CI) |  |
| --- | --- | --- | --- |
| **Fecundity** | Outbred | 1.05 (0.67-1.64) |  |
| **Hatching time in cold** | Outbred pilot | 7.07 (3.37-14.83) |  |
|  | Outbred replica 1 | 2.21 (1.49-3.26) |  |
|  |  | Males | Females |
| **CCRT** | Outbred replica 1 | 3.44 (2.31-5.18) | 1.89 (1.30-2.74) |
|  | Outbred replica 2 | 3.79 (2.54-5.67) | 5.18 (3.43-7.82) |
|  | Introgressed | 2.44 (1.64-3.62) | 4.16 (2.69-6.41) |
|  | Individual DGRP | 11.63 (6.79-19.93) | 2.26 (1.54-3.33) |
| **Survival after chill-coma** | Outbred | N/A | 7.80 (3.27-18.60) |
|  | Introgressed | N/A | 1.89 (0.99-3.62) |
|  | Individual DGRP | 9.94 (5.49-18) | 6.88 (3.43-13.82) |
| **Osmotic stress** | Outbred | N/A | 1.61 (1.21-2.13) |
| **Starvation stress** | Outbred | 1.52 (1.15-2.01) | N/A |
| **Developmental time** | Outbred pilot | 5.69 (2.72-11.94) |  |
|  | Outbred replica 1 | 2.62 (1.88-3.66) |  |
|  | Outbred replica 2 | 2.60 (1.94-5.88) |  |
|  | Individual DGRP | 1.95 (1.30-2.92) |  |

**Figure 1.**
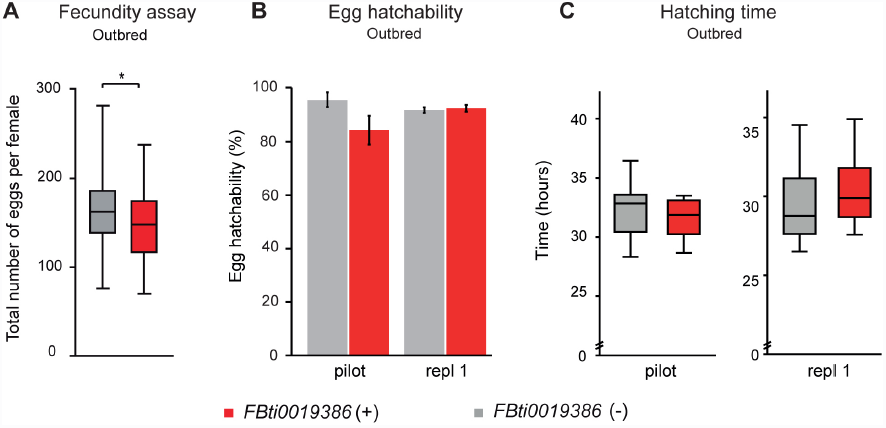
*FBti0019386* does not affect fecundity (A), egg hatchability (B) or hatching time (C) in outbred populations. (A) Average number of eggs laid by outbred females without *FBti0019386* insertion (grey) and with *FBti0019386* insertion (red). (B) Percentage of hatched embryos. (C) Average hatching time. In all cases, error bars represent standard error of the mean (S.E.M.).

We then checked whether outbred flies with and without *FBti0019386* differed in egg hatchability and/or hatching time. We first performed a pilot experiment using 150 embryos per strain and we found that flies with the insertion did not show significant differences compared to flies without the insertion in egg hatchability (t test, p-value > 0.05) (Figure 1B) or hatching time (t test, p-value > 0.05) (Figure 1C). Although differences were not significant, flies with the insertion showed a lower number of hatched eggs (Figure 1B) and a shorter hatching time (Figure 1C). We thus repeated the experiments using 500 embryos per strain and we found that flies with and without *FBti0019386* did not differ in egg hatchability (t test, p-value > 0.05) or hatching time (t test, p-value > 0.05).

Overall, we did not find significant differences in fecundity, egg hatchability or egg hatching time in flies with and without *FBti0019386* insertion. These results suggest that *FBti0019386* does not have a significant effect on these phenotypes.

### *FBti0019386* insertion does not affect egg hatching or egg-to-adult viability under cold stress conditions

As mentioned above, *Bin1* plays a role in general environmental stress response in *Drosophila* (Costa, et al. 2011). We thus screened several phenotypes in embryos under cold stress conditions: egg hatching, egg hatching time, and egg-to-adult viability.

**Figure 2.**
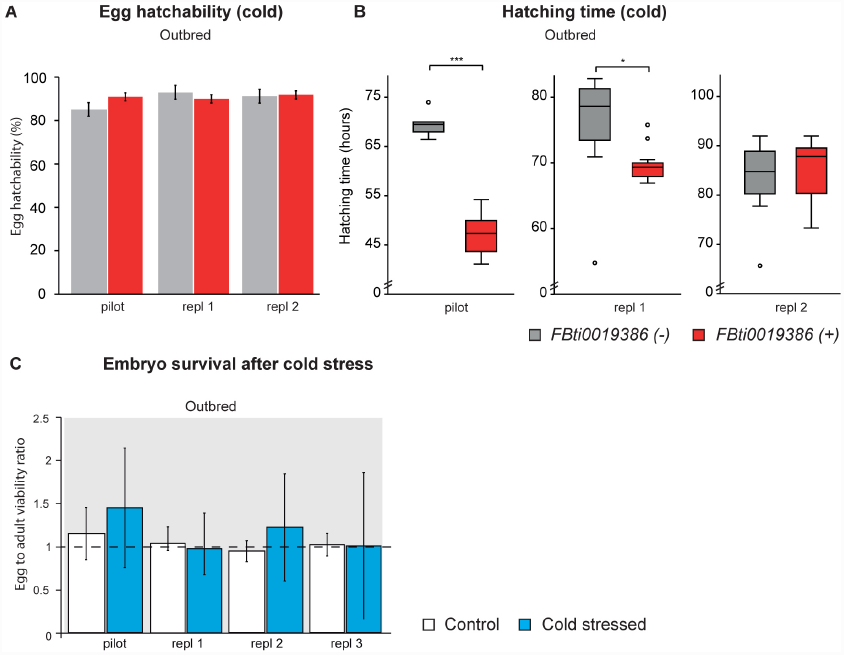
*FBti0019386* does not affect embryo hatching or survival in cold stress conditions in outbred populations. (A) Percentage of embryos that hatched during cold-stress periods (see Material and Methods). (B) Average egg hatching time. (C) Egg-to-adult survival after a single cold stress period during embryonic stage. Bars represent the survival ratio between flies with *FBti0019386* and flies without *FBti0019386* and error bars represent S.E.M.

We performed egg hatching and egg-hatching time assays in outbred populations under repeated cold stress exposure (see Material and Methods). We did not detect differences in egg hatching between flies with and without the insertion in any of the three replicas performed (t test, p-value > 0.05) (Figure 2A). However, flies with *FBti0019386* insertion from the pilot experiment and the first replica hatched significantly before flies without the element (t test, p-value << 0.001 and p-value = 0.011, respectively) (Table 2) while no differences were observed in the second replica (t test, p-value > 0.05) (Figure 2B).

We further tested whether flies with and without *FBti0019386* differed in the egg-to-adult viability after exposing outbred flies to a single cold-stress period during early embryo stages. Our results showed that there are no differences in survival between flies with and without the insertion in control conditions or under cold-stress (two-way ANOVA, p > 0.05, Figure 2C).

Overall, and although variability in hatching time was observed in some of the experiments performed, our results suggest that *FBti0019386* insertion does not affect cold-tolerance during the embryo stage.

### *FBti0019386* is associated with increased sensitivity to cold stress in adults

Because we could not find any significant difference between strains with and without *FBti0019386* in embryonic stage, we decided to test whether differences between the two strains were present in adult phenotypes. We first tested whether adult flies with and without *FBti0019386* insertion differed in chill-coma recovery time (CCRT) and survival after cold stress. CCRT is used as a reliable measure of cold tolerance in *Drosophila* (Gibert, et al. 2001; Macdonald, et al. 2004). We observed that flies with the insertion showed significantly longer recovery time compared to flies without the insertion suggesting that they were more sensitive to cold stress (Mann-Whitney test, p-value << 0.001) (Figure 3A and Table 2). We replicated this result in flies with the same genetic background (Mann-Whitney test, p-value < 0.05) and in flies with two other genetic backgrounds: the introgressed strains generated in our laboratory (Mann-Whitney test, p-value << 0.001) and a couple of inbred strains from the DGRP project (Mann-Whitney test, p-value << 0.001) (Figure 3A and Table 2) (see Material and Methods).

**Figure 3.**
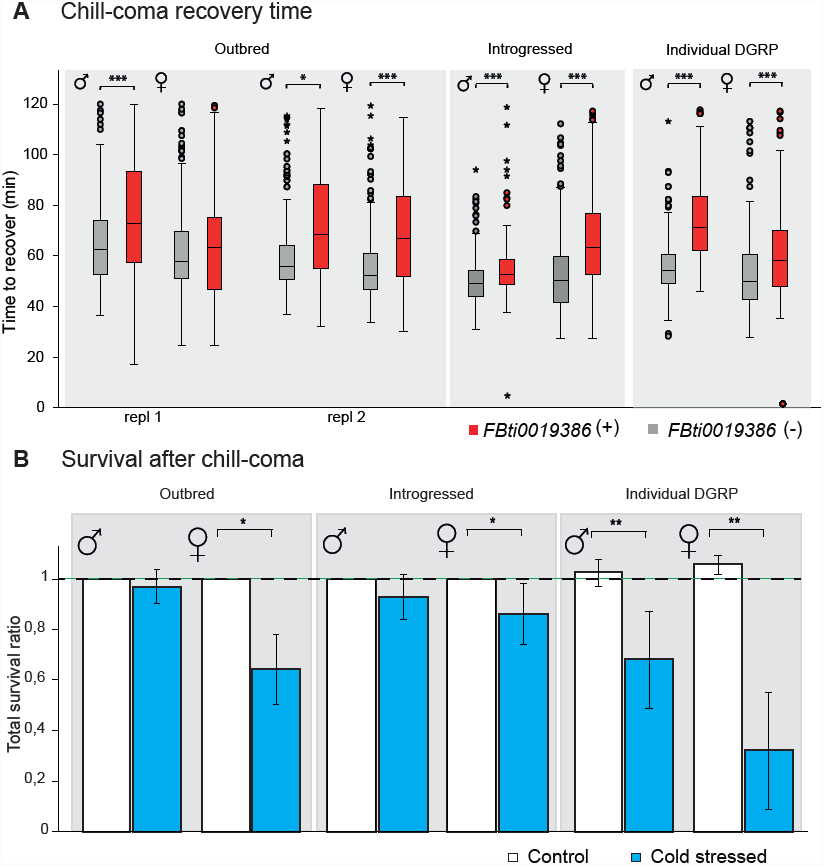
Flies with *FBti0019386* insertion are more sensitive to cold stress. (A) Average time to recover after chill coma in adult flies from outbred populations, introgressed strains and inbred DGRP strains (RAL-857 and RAL-802). (B) Survival ratio between flies with the element and flies without the element after chill coma exposure (blue) and in control conditions (white) in the three genetic backgrounds. Error bars represent S.E.M.

In accordance with this increased cold sensitivity, flies with the insertion also showed an increased mortality following chill-coma exposure, although these differences were not always significant (Figure 3B and Table 2).

Finally, we also tested whether flies with *FBti0019386* insertion were more sensitive to osmotic stress and starvation stress. We found that outbred females with the insertion were more sensitive to high salt concentrations (Kaplan-Meyer, log Rank p-value < 0.001) (Figure S3A and Table 2), and outbred males with the insertion were more sensitive to starvation stress (Kaplan Meyer, log rank p-value < 0.001) (Figure S3B and Table 2).

Overall, longer CCRT and lower cold-stress survival in flies with *FBti0019386* insertion across backgrounds suggested that this mutation is negatively affecting adult cold-stress response. This high sensitivity to cold stress likely represents the cost of selection of this TE mutation. Furthermore, preliminary results are suggestive but not conclusive of a negative role of *FBti0019386* in general response to stress.

### *FBti0019386* insertion is associated with shorter developmental time

During the course of the experiments, we noticed that flies with *FBti0019386* showed a shorter developmental time (DT) than flies without the insertion. Because DT is relevant to fitness in all organisms, and especially for those such as *D. melanogaster* that occupy ephemeral hábitats (Chippindale, et al. 1997), we decided to test this observation. We found that outbred flies (Mann-Whitney test, pilot experiment p-value = 0.006 and replica 1 and 2 p-value < 0.001) and inbred DGRP flies (t test, p-value = 0.02) with the insertion developed faster compared to flies without the TE insertion (Figure 4 and Table 2). On average, flies with *FBti0019386* insertion developed 9.4 to 17.9 hours before compared to flies without the insertion. However, we could not detect significant DT differences in the introgressed strains differing by the presence/absence of *FBti0019386* (t test, p-value > 0.05) (Figure 4), suggesting that polymorphisms other than the TE influence DT in this background.

**Figure 4.**
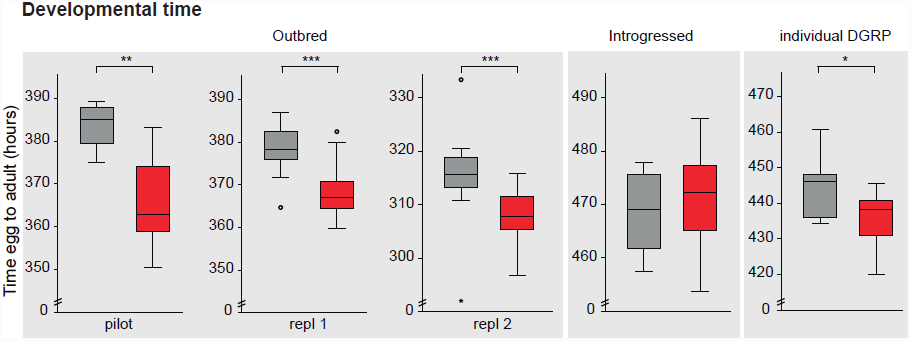
*FBti0019386* is associated with shorter developmental time. Average egg-to-adult developmental time in populations without *FBti0019386* (grey) and with the *FBti0019386* insertion (red). Error bars represent S.E.M.

### *FBti0019386* frequency showed clinal patterns in North America and Australia but not in Europe

Shorter DT and increased sensitivity to cold stress are not consistent with a role of *FBti0019386* in temperate adaptation (Gonzalez, et al. 2010). However, previous evidence for a role in temperate adaptation was based on the analysis of only two North American and five Australian populations (Gonzalez, et al. 2010). To further test these results, we estimate *FBti0019386* frequencies in additional populations from North America, Australia, Europe and Africa (Table S5) using *T-lex2* pipeline (Fiston-Lavier, et al. 2014). We found that *FBti0019386* insertion is present at 10% frequency in a Rwanda population confirming its low frequency in Africa (Table S5). We confirmed that the TE is present at intermediate to high frequencies in 20 additional out-of-Africa populations (Figure 5 and Table S5). We also confirmed that the TE frequency varies clinally with latitude in North America and Australia (Pearson correlation p-value = 0.008 and p-value= 0.002 respectively; Table S6).

**Figure 5.**
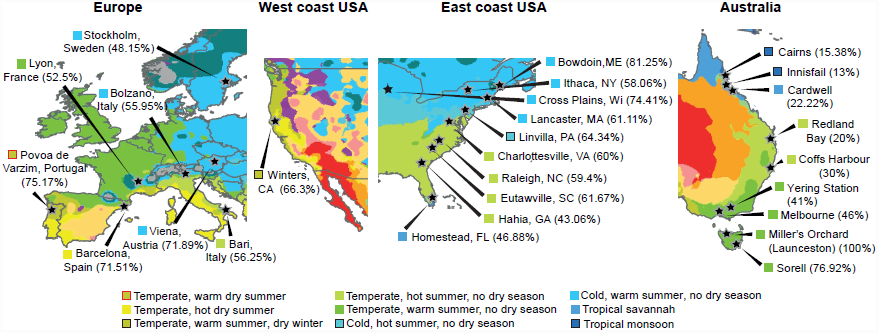
Climate map with *Drosophila melanogaster* population samples analyzed with Tlex. The frequency of *FBti0019386* for each population is shown in brackets. Climate maps are modified from Peel et al., 2007 (Peel, et al. 2007).

However, when we analyzed the *FBti0019386* frequency in seven European populations we did not find any significant latitudinal pattern (Pearson correlation p-value= 0.193; Table S6).

Besides latitude, we also tested whether other geographical and climatic variables showed significant correlations with *FBti0019386* frequency. We found significant correlations between frequency and temperature-related variables in North America and between frequency and both temperature-related and precipitation-related variables in Australia (Table S6). No significant correlation was found in Europe (Table S6). Because most of the climatic variables are significantly correlated among them and with latitude (Table S7), we performed a Principal Component Analysis (PCA) to disentangle the relationships between the variables. In North America, climate variables were grouped in four components, in Australia in three and in Europe in two (Table S8). As expected based on the correlation analyses, only in North America and in Australia, some of the climatic variables grouped with latitude and frequency (Figure S2A). In North America, the first component accounted for 45% of climatic variation (Table S9) and explained 33% of the variation in *FBti0019386* frequency (Figure S2). In Australia, the first component accounted for 68% of climatic variation (Table S9) and explained 89% of the frequency variation (Figure S2). Finally in Europe, the first principal component explained 62% of the climatic variation (Table S9) but was not significantly correlated with *FBti0019386* frequency (Figure S2).

Overall, although we were able to confirm the clinal pattern of *FBti0019386* in North America and Australia, our results indicate that not such pattern is present in Europe. In Australia, this clinal pattern is well explained by the observed climatic variation, while in North America climatic variation did not fully explain the observed correlation between *FBti0019386* frequency and latitude, suggesting that other factors might be involved in the observed clinal pattern. As expected, none of the climatic variables significantly correlated with TE frequency in Europe.

### *FBti0019386* is associated with down-regulation of sarah in female flies

To shed light on the molecular mechanism of *FBti0019386* insertion, we measured the expression of *sarah* and *Bin1* in non-stressed conditions in embryos and in non-stressed and cold-stress conditions in female flies with and without *FBti0019386* insertion.

We did not observe significant differences in *sarah* or *Bin1* expression in embryos differing by the presence/absence of *FBti0019386* insertion (t test, p-value > 0.05) (Figure 6A and 6B). However, we observed that adult female flies with the insertion showed a reduction of *sarah* expression compared to flies without the element both in control conditions and after cold-stress conditions, although results were only significant under control conditions (t test, p-value = 0.03) (Figure 6C). On the other hand, no significant differences in expression level were observed for *Bin1* (t-test, p-value > 0.05) (Figure 6D).

**Figure 6.**
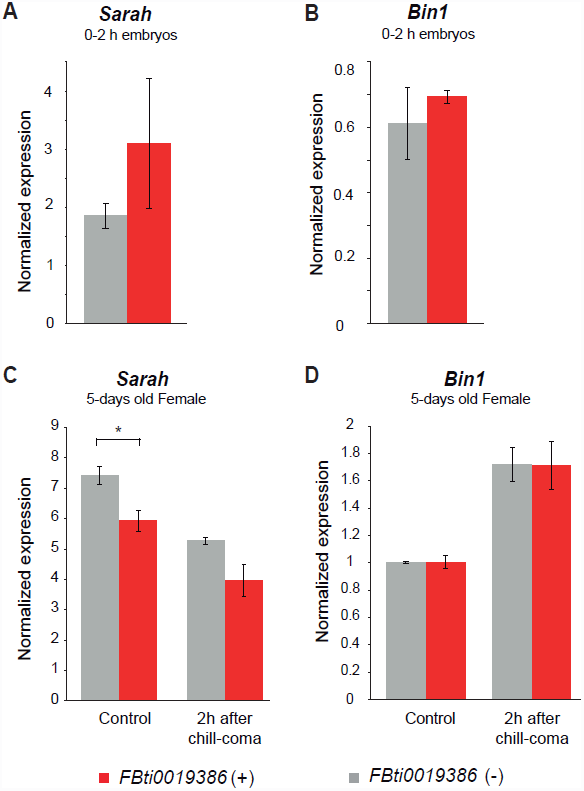
Flies with *FBit0019386* insertion showed *sra* down-regulation. Real-time PCR quantification of *sarah* and *Bin1* transcript levels in outbred flies without *FBti0019386* (grey bars) and with *FBti0019836* (red bars). We represent the average copy number of *sra* (A and C) and *Bin1* (B and D) relative to *Act5C* with S.E.M. error bars for three biological replicates in 0-2 hours embryos and in 5-days old females. Normalized expression measured 2 hours after chill-coma for *sra* and *Bin1* is depicted in C and D, respectively.

Interestingly, we observed a change in both *sarah* an *Bin1* expression after cold stress in flies with and without *FBti0019386* insertion: *sarah* is down-regulated in cold stress conditions (t test, p-value < 0.05 in both cases) while *Bin1* is up-regulated (t test, p-value < 0.05 in both cases).

Overall, we did not observed any change in expression of *sra* and *Bin1* in embryos, in agreement with the lack of phenotypic consequences of *FBti0019386* in this developmental stage. However, we observed a down-regulation of *sarah* in flies with *FBti0019386* insertion that was significant under non-stress conditions. Moreover, we showed that both *sarah* and *Bin1* changed their expression in response to cold stress.

### *FBti0019386* could be affecting sarah expression by ectopically assembling heterochromatin

TEs from the *invader4* family contain sites with homology to PIWI interacting RNAs (piRNAs) that act as *cis*-acting targets for heterochromatin assembly by recruiting Heterochromatin Protein 1a (HP1a) (Sentmanat and Elgin 2012). Specifically, these piRNA binding sites are located in the LTR sequences. Because *FBti0019386* is a solo-LTR, we hypothesized that ectopic assembly of heterochromatin induced by *FBti0019386* insertion might be the mechanism underlying *sra* down-regulation. We first checked whether there is evidence for piRNAs binding to *FBti0019386* sequence and we found that both sense and antisense piRNAs can bind to this TE (Figure 7A) (see Material and Methods). Second, we tested whether there is evidence for the presence of HP1a binding to *FBti0019386* sequence. We analyzed the region containing *Bin1*, *sra*, and *FBti0019386* and found that HP1a specifically binds to *FBti0019386* sequence (Figure 7B) (see Material and Methods). Thus, these results suggest that down-regulation of *sra* in flies with the insertion could be due to the piRNA-mediated heterochromatin assembly induced by *FBti0019386*.

**Figure 7.**
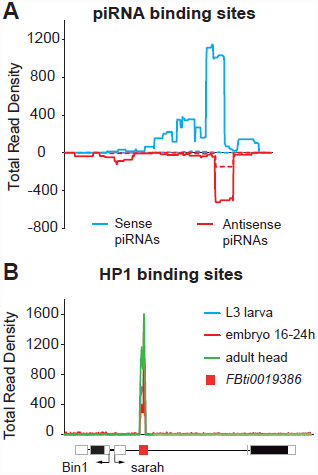
*FBti0019386* could bind piRNA and HP1 protein. (A) Mapping of piRNA sense (blue) and anti-sense (red) RNA-seq reads against *FBti0019386* sequence. Data from Li et al., 2009 is depicted in dashed lines and data from Satyaki et al., 2014 is represented in continuous line. (B) Mapping of reads coming from HP1a ChIP-Seq experimental data against the genome region containing *Bin1*, *FBti0019386* and *sra*. Experimental data from L3 larva (blue), 16-24h embryo (red) and adult head (green) is given.

## DISCUSSION

In this work, we explored the plausible phenotypic space of the putatively adaptive *FBti0019386* insertion in different developmental stages, embryo and adult, and in different environmental conditions, non-stressed and cold, osmotic, and starvation stress. Overall, we found that *FBti0019386* mediates sensitivity to cold stress conditions (Figure 3) and is associated with faster developmental time (Figure 4). These two phenotypic effects have plausible fitness consequences in nature that could explain why the mutation increased in frequency in natural populations but has not reach fixation. Increased sensitivity to cold stress conditions is likely to reduce fitness of the flies that carry *FBti0019386* insertion, and likely represents the cost of selection of this mutation. On the other hand, faster developmental time is likely to increase the fitness of flies with *FBti0019386* insertion. In nature, quick development favors *D. melanogaster* individuals for several reasons. First, larvae feed on rotting fruits that are ephemeral, so quick development allows them to pupate before the food source is exhausted. Second, competition increases as more and more eggs are laid on a piece of fruit, also favoring individuals with faster developmental time (Nunney 1990). Third, breeding sites in nature can be destroyed by physical factors and predation, individuals that develop faster are thus more likely to escape microhabitat destruction. And fourth, faster developmental time accelerates the age of first breeding, which is relevant for the organism if most reproduction happens in expanding populations. This is the case for *D. melanogaster* populations that expand their population size every spring. Thus, it is plausible that *FBti0019386* increased in frequency in natural populations because of its positive effect on developmental time while it did not reach fixation because of its negative effect on cold-stress resistance. Our results emphasize the importance of exploring different phenotypes to fully characterize the effects of natural mutations, as have been suggested before (Mackay 2010; Guio, et al. 2014).

By combining several tests that capture different signatures of selection at the DNA level, we demonstrate that *FBti0019386* shows signatures of positive selection suggesting that it is an adaptive mutation (Table 1). However, our results also suggest that *FBti0019386* is not a clinal mutation likely to be involved in temperate adaptation as has been previously proposed (Gonzalez, et al. 2010). First, adaptation to temperate climates has been associated with increased stress resistance, increased developmental time and decreased fecundity (Stanley and Parsons 1981; Hoffmann, et al. 2003; Schmidt, et al. 2005; Folguera, et al. 2008; Schmidt and Paaby 2008) but see also (James and Partridge 1995; James, et al. 1997; Trotta, et al. 2006). However, we showed that *FBti0019386* is associated with increased sensitivity to cold stress (Figure 3), with shorter DT (Figure 4) and does not significantly affect fecundity (Figure 1). Thus, the phenotypic effects of *FBti0019386* are not consistent with a role of this insertion in temperate adaptation. Second, our global analyses of *FBti0019386* population frequency showed that *FBti0019386* frequency correlates with latitude and with climatic variables in North America and in Australia but not in Europe (Figure 5, Table S6). We suggest that the clinal frequency patterns in North America and in Australia could be due to the dual colonization of these two continents by European and African populations rather than to the operation of spatially varying selection (Caracristi and Schlotterer 2003; Rouault, et al. 2004; Duchen, et al. 2013; Bergland, et al. 2014). The lack of clinal frequency patterns in Europe supports this conclusion. Thus, genome-wide scans that identify loci that are differentiated between populations should be taken as a first step towards the identification of loci that are subject to spatially varying selection (Gonzalez, et al. 2010; Kolaczkowski, et al. 2011; Fabian, et al. 2012; Reinhardt, et al. 2014). Further functional validation should be gathered before concluding that the candidate loci are under spatially varying selection (Bergland, et al. 2014).

Our results also shed light on the molecular processes that lead from genotype to phenotype. We found that *FBti0019386* is associated with down-regulation of *sra* (Figure 6C), most probably by inducing the ectopic assembly of heterochromatin by recruiting HP1a protein. As previously described for other elements from the *invader4* family, we showed that *FBti0019386* has piRNA binding sites (Figure 7A) (Sentmanat and Elgin 2012). We also showed that HP1a binds specifically to *FBti0019386* sequence, further suggesting that down-regulation of *sra* is the result of piRNA-mediated heterochromatin assembly. These results highlight the role of TE remnants as silencing signals to be used by piRNAs to direct heterochromatin formation (Sentmanat, et al. 2013).

A recent update of FlyBase, the database of Drosophila genes and genomes, annotated two new *Bin1* transcripts that have their transcription start site inside *FBti0019386* (St Pierre, et al. 2014). As a consequence, these two new transcripts would only be produced in strains with the insertion, and could contribute to differences in the level of *Bin1* expression in flies with and without the insertion. Although we did not find differences in *Bin1* expression, we cannot discard that differences in the level of expression of *Bin1* are present in developmental stages, tissues, or environmental conditions that we have not investigated.

Although *sra* and *Bin1* have not been associated with developmental time, both genes play important roles during development and have been associated with a wide range of processes (Chang, et al. 2003; Ejima, et al. 2004; Horner, et al. 2006; Takeo, et al. 2006; Chang and Min 2009; Matyash, et al. 2009; Takeo, et al. 2010; Costa, et al. 2011; Nakai, et al. 2011). A genome-wide screening looking for genes influencing DT in *D. melanogaster* has shown that the many candidate genes were involved in a wide range of biological processes such as cellular metabolic processes, organismal development, and response to stress (Mensch, et al. 2008). More recently, developmental timing in insects has been associated with hormonal and circadian control (Di Cara and King-Jones 2013; Yadav, et al. 2014). Interestingly, *sra* is regulated by *Shaggy*/*GSK-3*β *(sgg)*, a Ser-Thr kinase involved in the regulation of circadian rhythmicity (Martinek, et al. 2001). On the other hand, both *Bin1* and *sra* are stress-response genes: *Bin1* is up-regulated in response to stress and *sra* is down-regulated (Figure 6). *Bin1* is a known key player in transcriptional response to environmental stress (Costa, et al. 2011). While there was no previous evidence for a direct role of *sra* in response to stress, *sra* could be affecting stress response through its role in the calcium pathway (Takeuchi, et al. 2009; Teets, et al. 2013; Davies, et al. 2014). *sra* inhibits *calcineurin*, a highly conserved protein in eukaryotes that has the ability to sense calcium (Hogan, et al. 2003). Although it is not deeply understood, calcium pathways play a role during general cell-stress response including cold stress response (Takeuchi, et al. 2009; Teets, et al. 2013; Davies, et al. 2014). Note that many genes that affect complex traits in Drosophila had well-characterized roles in early development and were not previously annotated to affect adult quantitative traits (Mackay 2010).

*FBti0019386* adds to the growing list of TE-induced adaptive mutations that have been linked to their fitness effects and their underlying molecular mechanisms (Schmidt, et al. 2010; Magwire, et al. 2011; Guio, et al. 2014; Mateo, et al. 2014; Sun, et al. 2014). Overall, these examples highlight the variety of mechanisms underlying adaptive mutations and point towards a significant role of TEs in response to stress. However, the number of characterized mutations is still too small to obtain an overall picture of adaptation. In depth characterization of a representative set of adaptive mutations in natural populations will allow us to start answering long-standing questions such as which traits are more relevant for adaptation? What is the effect-size distribution of adaptive mutations?, and What evolutionary processes underlie adaptive evolution?

## MATERIAL AND METHODS

### Sequence analysis of the *FBti0019386* flanking regions

SNP data was downloaded form the DGRP2 webpage (https://www.hgsc.bcm.edu/arthropods/drosophila-genetic-reference-panel) in *vcf* format. Strains with (N=65) and without (N=38) *FBti0019386* insertion were filtered using *vcftools v_0.1.10 (http://vcftools.sourceforge.net/).* Nucleotide diversity (π), Tajima´s D, and the CL (Composite Likelihood of SNPs, Nielsen et al. 2005) statistic, were calculated for the two sets of sequences using the PopGenome package in R (Pfeifer, et al. 2014). Sliding windows analyses were performed for 200bp size windows spanning 1kb and 2kb regions flanking the insertion. Differences between strains with and without the insertion were more drastic for the 1kb region flanking the insertion, therefore we focused our analysis in this region.

Simulations were performed using the MS program (Hudson 2002). Theta values were estimated using the 205 DGRP2 strains for the 2kb region around *FBti0019386* (theta = 4.77/kb) and for the 3R chromosomal arm (theta = 4.5/kb). Thus, simulations were performed for theta values of 4/kb and 5/kb, which are frequently used as neutral values in *D. melanogaster*.

*Ad hoc* perl scripts were used for the re-sampling analyses. 1,000 random samples of the 103 DGRP strains analyzed were obtained keeping the same proportion as in the original present and absent datasets (60%/40%, respectively) and a sample size of nearly the 50% of the total dataset.

We also computed CLR (Composite Likelihood Ratio) as 2*(logCL (present) –logCL (absent)), for a 1kb region around the TE insertion. Because demography could produce similar patterns as positive selection, we performed a random sampling of 1,000 1kb-long regions from the 3R chromosome for the absent and present datasets and calculated nucleotide diversity, Tajima´s D, CL, and CLR statistics in each one of them.

### Fly stocks

*Outbred strains.* We selected six inbred strains from the Drosophila Genetic Reference Panel (Mackay, et al. 2012; Huang, et al. 2014) homozygous for the presence of *FBti0019386* insertion (RAL-21, RAL-40, RAL-177, RAL-402, RAL-405 and RAL-857). We placed 10 virgin females and 10 virgin males of each strain in a fly chamber to create an outbred population sharing the TE insertion. We also selected six inbred strains without the insertion (RAL-75, RAL-138, RAL-383, RAL-461, RAL-822 and RAL-908) and created an outbred strain following the same procedure explained above. Each outbred population was maintained by random mating (N ≈ 800 flies per generation) for at least 10 generations before starting the experiments.

*Introgressed strains*. We selected two DGRP strains: one homozygous for the presence of *FBti0019386* insertion (RAL-177) and one homozygous for the absence (RAL-802). We crossed RAL-177 virgin females with RAL-802 males and backcrossed the virgin females from the following generations with RAL-802 males for 12 generations. After that, we did brother-sister crosses until we obtained homozygous strains for the absence and homozygous strains for the presence of *FBti0019386*.

*Individual DGRP strains.* We used a couple of individual DGRP strains differing by the presence/absence of the insertion to perform our phenotypic assays. We used RAL-857 (homozygous for the presence *FBti0019386*) and RAL-802 (homozygous for the absence).

### Phenotypic assays

All experiments were performed using outbred populations. Additionally, we used introgressed and individual DGRP strains to perform chill-coma recovery time assay, survival after chill-coma, and developmental time assays.

#### Fecundity

40 virgin females from each strain were placed individually in vials with one male from the same strain. During 17 days flies were moved to new vials every two days and the number of eggs laid per female during that period was counted. Average of the total number of eggs laid per female was compared between flies with and without *FBti0019386*.

#### Egg hatchability and hatching time

4 to 8 day-old flies were allowed to lay eggs for 3 hours on apple juice – agar medium with fresh yeast. Embryos were separated in groups of 20 or 50 and placed into food vials. Vials were kept at room temperature (19 ºC-22 ºC) and checked during the following hours for hatched eggs (2-5 times per day). We analyzed the estimated average time over the midpoint of each successive interval in order to estimate the hatching time. Two experiments were performed following this protocol: a first pilot experiment with 150 embryos per strain, and one replica with 500 embryos per strain.

Egg hatchability and egg hatching time was also analyzed under cold stress conditions. Embryos were placed at 1ºC overnight for 14 hours and at 18ºC during the day, and this cycle was maintained until all the eggs had hatched. We performed a pilot experiment with 100 embryos per strain and additional experiments with 240 and 160 embryos per strain respectively.

#### Cold stress in embryos

7 to 10 day-old flies were allowed to lay eggs for 3 hours on apple juice-agar medium with fresh yeast. Embryos were collected following the methodology described in Schou (2013), and placed into food vials in groups of 50. When embryos were 3 to 6 hours old, vials were placed at 1ºC for 14 hours, and maintained at 18ºC until adult emergence. Simultaneously, control vials were always maintained at 18ºC and not cold-exposed to control for other variables affecting egg to adult survival. We performed a first pilot experiment using 280 embryos per strain and three biological replicas using 350 embryos per strain (in replica 1) and 750 embryos per strain (in replicas2, and in replica 3). In all cases, we analyzed egg to adult survival after all the adults had emerged.

#### Chill-coma recovery time

3 to 5 day-old flies were separated by sex and by strain and placed into 5 empty vials in groups of 50. We allowed flies to recover from CO_2_ anaesthesia for 1 hour and then, vials were put in ice and kept in a 4ºC chamber for 16 hours as described in David et al., (1998). After the cold shock, adults were transferred to Petri dishes at room temperature (22ºC- 24ºC), and recovery time was monitored for successive intervals of 30 seconds during 2 hours. We considered as recovered flies those that were able to stand on their legs. As a control, we monitored survival of flies that were kept at room temperature: 3 vials of 20 flies each, by sex and strain.

#### Survival after chill-coma

5 to 8 day-old flies were separated by sex and strain and placed into 6 food vials in groups of 20. We allowed flies to recover from CO_2_ anaesthesia for at least 2 days. After that, flies were changed to empty food vials and were put in ice, and kept in a 4ºC chamber for 16 hours. When adults were recovered from chill-coma, we transferred them to food vials and we monitored mortality during the next 5 days. As a control, we monitored survival of flies that were kept at room temperature: 3 vials of 20 flies each, by sex and strain.

#### Osmotic stress

4 to 7 day-old flies were separated by sex and strain and placed in groups of 20 into 20 food vials containing 3% of NaCl, and into 5 vials with normal food as a control. Flies were maintained at room temperature (22ºC- 24ºC) and dead flies were counted every 12-24 hours until all the treated flies were dead.

#### Starvation stress

3 to 4 day-old flies per strain were separated by sex and strain and placed in groups of 20 into 20 food vials containing only 1.5% agar, and into 5 vials with normal food as a control. Flies were maintained at room temperature (22ºC- 24ºC) and dead flies were counted 3 times a day until all the treated flies were dead.

#### Developmental time

7 to 10 day-old flies were allowed to lay eggs for 3 hours. A total of 500 embryos per strain were collected and distributed in groups of 50 per food vial and were maintained at 18ºC. Vials were checked every 6-8 hours for emerging adults until all flies had emerged. We estimated the average DT over the midpoint of each successive interval.

### Statistical analyses

Analyses were performed with SPSS v21. We first tested if data followed a normal distribution by performing Kolmogorov-Smirnov test. t-test was performed for normal data and Mann-Whitney for non-normal data. Survival curves were compared with log-rank test. When statistical test were significant, we estimated the size effect of the mutation by calculating the odds-ratio and its confidence interval.

### *FBti0019386* frequency estimation for natural populations

To obtain *FBti0019386* frequency, we run *T-lex2* (Fiston-Lavier, et al. 2014) using *Drosophila* whole-genome sequences available from a total of 23 populations from North America, Australia, Europe, and Africa (Table S5).

The accuracy of TE frequency estimates using *T-lex2* is affected by coverage. However, coverage for all samples was higher than 20x except for Lyon (France) and California (USA), which had 8x and 4.7x coverage respectively, suggesting that overall frequency estimates are accurate.

### Correlation analysis of FBti0019386 frequency with geographic and climate variables

We analyzed whether the frequency of *FBti0019386* insertion correlated with different geographical and climatic variables in North America, Australia and Europe using Pearson product-moment correlations. We also performed a principal component analysis (PCA) to disentangle the relationships between the climatic variables using Statistica (v8.0, StatSoft, Inc. 2007). Climatic data was obtained from the weather stations adjacent to collection sites of each population, available in Peel et al. (2007). When necessary, data were transformed as described in Sokal and Rohlf (1995) (see pages 411-422).

### mRNA transcript levels analysis (qRT-PCR)

Total RNA was extracted from three biological samples of 40 adult females (5 day-old) from outbred populations differing by the presence/absence of *FBti0019386* insertion using Trizol reagent and PureLink RNA Mini kit (Ambion). RNA was treated on-column with DNase I (Trizol) and after RNA purification. Reverse transcription was carried out using 1 µg of total RNA, Anchored-oligo(dT) primer and Transcription First Strand cDNA Synthesis Kit (Roche). The resulting cDNA was used for qRT-PCR with SYBR Green (BioRad) on an iQ5 Thermal cycler. *sarah* total expression was measured using a pair of primers specific to a 124 bp cDNA amplicon spanning the 5’UTR/exon junction of the gene (5’-ACAACAACGGTGGAGAAGAGCCGT-3’ and 5’-GGTGCATCGGCGGACGCATTG-3’). For *Bin1*, we measured the 66 bp cDNA amplicon spanning the 5’UTR/exon junction using specific primers (5’-TGTCGTCCCGTAGAGCAGAA-3’ and 5’-CAAGCAGATTGACCGCGAGA-3’). In both cases, we normalized the expression with *Act5C* (5’-GCGCCCTTACTCTTTCACCA-3’and 5’-ATGTCACGGACGATTTCACG-3’). Expression was measured in non-stressed conditions and in cold-stress conditions: 16 hours at 4 ºC and 2 hours at room temperature to allow flies to recover.

We also analysed the expression of both genes in 0-2 hours embryos using the same procedure. We collected the embryos from population cages containing approximately 800 flies from outbred populations differing by the presence/absence of *FBti0019386* insertion. Briefly, 4 to 8 day-old flies were allowed to lay eggs for 2 hours on apple juice - agar medium with fresh yeast. Then, embryos were collected using a small brush and cleaned with water. Embryos were dechorionized by submerging them for 5 minutes in 50% bleach. After that, embryos were placed in a microcentrifuge tube, the excess of water was eliminated and the samples were freezed at -80 ºC until RNA extraction.

### Detection of piRNA reads binding to *FBti0019386* sequence

We used small RNA sequencing data to check whether piRNAs reads mapped to *FBti0019386* sequence, following a methodology similar to that described in Sentmanat and Elgin (2012). Briefly, we obtained the small RNA reads from Oregon R ovaries (accession number SRP000458) (Li, et al. 2009), and from wild type ovaries (accession number: SRX470700) (Satyaki, et al. 2014). We aligned the reads to *FBti0019386* sequence by using BWA-MEM package version 0.7.5a-r405 (Li 2013). Then, we used samtools and bamtools (Barnett, et al. 2011) to index and filter by sense/antisense reads. Finally, we obtained the total read density using R (Rstudio 2012).

### Detection of HP1a protein binding in *FBti0019386* sequence

We downloaded all available raw data from modEncode HP1a protein ChIP-Seq experiments: embryos (ID 3391 and 3392), 3rd instar larvae (ID 4936) and adult heads (ID 5592) (http://data.modencode.org). Then, we mapped the reads against the sequence obtained from *Drosophila* reference genome, containing *Bin1* and *sra* genes, and *FBti0019386*, corresponding to the chromosomal coordinates 3R: 12,010,721-12,025,306 (release 5). We performed the alignments following the same methodology as for the piRNA reads analysis.

## ACKNOWLEDGEMENTS

We thank Lain Guio and Miriam Merenciano for comments on the manuscript. AU is a FPI fellow (BES-2012-052999) and JG is a *Ramón y Cajal* fellow (RYC-2010-07306). This work was supported by grants from the European Comission (Marie Curie CIG PCIG-2011-293860) and from the Spanish Government (Fundamental Research Projects Grant BFU-2011-24397) to J.G.

